# A Telescopic Independent Component Analysis on Functional Magnetic Resonance Imaging Data Set

**DOI:** 10.1101/2024.02.19.581086

**Authors:** Shiva Mirzaeian, Ashkan Faghiri, Vince D. Calhoun, Armin Iraji

## Abstract

Brain function can be modeled as the dynamic interactions between functional sources at different spatial scales, and each spatial scale can contain its functional sources with unique information, thus using a single scale may provide an incomplete view of brain function. This paper introduces a novel approach, termed “telescopic independent component analysis (TICA),” designed to construct spatial functional hierarchies and estimate functional sources across multiple spatial scales using fMRI data. The method employs a recursive ICA strategy, leveraging information from a larger network to guide the extraction of information about smaller networks. We apply our model to the default mode network (DMN), visual network (VN), and right frontoparietal network (RFPN). We investigate further on DMN by evaluating the difference between healthy people and individuals with schizophrenia. We show that the TICA approach can detect the spatial hierarchy of DMN, VS, and RFPN. In addition, TICA revealed DMN-associated group differences between cohorts that may not be captured if we focus on a single-scale ICA. In sum, our proposed approach represents a promising new tool for studying functional sources.

**Author summary:** Our study introduces “telescopic independent component analysis (TICA)”, a novel approach using a recursive ICA strategy for exploring brain function across multiple spatial scales using fMRI data. TICA constructs spatial hierarchies and identifies functional sources such as default mode network (DMN), visual network (VN), and right frontoparietal network (RFPN). By applying TICA, we uncovered hierarchical structures within these networks and revealed differences in the DMN between healthy individuals and those with schizophrenia. This approach offers a promising new tool for studying function dynamics comprehensively.

## 1. Introduction

### 1.1. Intrinsic connectivity networks

A Macroscopic brain’s function can be conceptualized as coordination and interaction among functional sources, a model incorporating the fundamental principles of segregation and interaction (Genon et al., 2018; Armin Iraji et al., 2022). From this perspective, the brain is comprised of distinct functional sources that dynamically interact with each other. These functional sources can be characterized by temporally synchronized patterns, which can be assessed at the macroscale using resting-state functional magnetic resonance imaging (rs-fMRI) and techniques such as independent component analysis (ICA)(Calhoun & Adalı, 2012; van den Heuvel & Hulshoff Pol, 2010). ICA is a robust blind source separation technique that separates complex rs-fMRI signals into spatial patterns that are temporally coherent and maximally independent of each other (Calhoun & Adalı, 2012; Calhoun et al., 2001; Calhoun & de Lacy, 2017). These spatial patterns and associated time courses formed the estimation of functional sources and are commonly referred to as intrinsic connectivity networks (ICNs) (Allen et al., 2014). Importantly, these ICNs can be captured at different spatial scales (Abou-Elseoud et al., 2010; Armin Iraji et al., 2022; Iraji et al., 2023). Previous research has underscored the significance of the analysis of ICNs at multiple spatial scales (Armin Iraji et al., 2022; Li et al., 2018; Meng et al., 2022; Meng et al., 2023). Previous studies that explore multiple spatial scales can be divided into two major categories. The category relies on a singular set of nodes, such as predefined regions or a single model-order ICA, to reconstruct different scales of functional hierarchy using various modularity or clustering techniques (Doucet et al., 2011); however, these approaches have no information that can be explained by looking at the other spatial scales. The other category of approaches appreciates that ICNs at different spatial scales can contain distinctive functional sources with their unique information. For instance, previous studies utilized ICA with different model orders to capture ICNs at different spatial scales to study whole-brain functional connectivity (Armin Iraji et al., 2022; Meng et al., 2022). Lower model order yields large-scale, spatial distributed ICAs (Beckmann et al., 2005; Calhoun et al., 2008; Iraji et al., 2016) while higher model orders result in more finely-grained, spatially specific ICNs (Abou-Elseoud et al., 2010; E. Allen et al., 2011). Importantly, the interaction between large-grained scale and small-grained scale can convey crucial insights into the brain’s functioning. for instance, multiscale brain analysis better captures sex-specific schizophrenia changes (Armin Iraji et al., 2022). However, these techniques require post hoc analysis to construct the spatial hierarchy and assign ICNs to different spatial scales, as the method does not impose direct constraints on the size of ICNs.

Here, we present a new ICA strategy, called Telescopic ICA (TICA), to construct spatial functional hierarchy and estimate ICNs across multiple spatial scales. The approach is a recursive technique that leverages ICNs from the previous scale to guide the ICA to obtain ICNs for another scale. Notably, our approach is designed to show how a large-scale network decomposes through a small-scale network using zooming in based on the prior level of the functional source.

### 1.2. Schizophrenia

Schizophrenia represents a psychotic disorder characterized by different cognitive impairments and a decline in both social and occupational functioning(Association, 2013; Van Den Heuvel & Fornito, 2014). This condition is multifaceted, diagnosed through a syndromic approach, and involves excluding other potential diagnoses. Schizophrenia lacks distinctive symptoms and is primarily diagnosed based on clinical observations of positive symptoms (such as delusions, hallucinations, disorganized speech, and disorganized or catatonic behavior) as well as negative symptoms (including apathy, blunted affect, and anhedonia), along with a noticeable deterioration in social functioning(Association, 2013). Schizophrenia exhibits substantial overlap with both schizoaffective disorder and psychotic bipolar disorder, in both symptoms, genetic factors, and other biomarkers(Clementz et al., 2016). Schizophrenia is theorized to be a developmental disorder marked by disrupted brain function, characterized by either functional dysconnectivity or alterations in functional integration(Friston & Frith, 1995; Kahn et al., 2015). research also has elucidated sex differences in the incidence and clinical manifestation of mental disorders (Dalsgaard et al., 2020; Paus et al., 2008). Females with schizophrenia tend to show more depressive symptoms, whereas males often experience more negative symptoms (Ochoa et al., 2012). Also, recent studies highlight the importance of analyzing the brain in multiscale to capture sex-specific schizophrenia changes (Armin Iraji et al., 2022); and previous studies have shown high-order functional connectivity applied to the diagnosis of psychotic disorders (Herzog et al., 2022; Li et al., 2023). Therefore, studying brain functional sources at different scales can provide crucial information about brain functional integration and its schizophrenia changes, potentially improving our understanding of the actual brain pathology underlying different schizophrenia subcategories. Here we used the TICA approach to identify the spatial map of the default mode network (DMN) in two scales and provide important information about brain functional sources and its schizophrenia changes, potentially improving our understanding of the actual brain pathology, thus eventually improving treatments and care for individuals with schizophrenia. Our results show that the TICA approach can capture significant group differences in DMN area between healthy people (HC) and schizophrenia patients (SZ) which are missed in single-scale ICA.

## 2. Materials and methods

### 2.1. Proposed approach

In our proposed approach, first, we applied two levels of principle component analysis (PCA) followed by group level ICA (GICA) on a data set (Calhoun et al., 2009; Erhardt et al., 2011). Subject spatial PCA at the first level applied to normalize the data and to allow subjects to contribute similarly to the common subspace. It also has denoising and computational benefits using the equation:

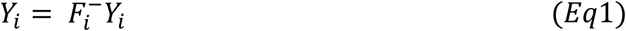

Here subject-specific BOLD signal *Y*_*i*_ is *T*_1_-by-*V* reduced data for a subject *i*, 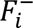 is the *T*_1_-by-*T* standardized reducing matrix.*T*_1_is the number of PCs retained for each subject, T is total number of time points and V is the number of voxels. While the subject-specific PCA privileges subject differences at the subject level, the group-level PCA favors subject commonalities(Erhardt et al., 2011). All subject-level principal components were concatenated together across the time dimension, and group-level spatial PCA was applied to concatenated subject-level principal components(*Eq2*, *Eq3*).

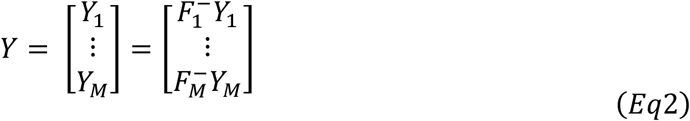

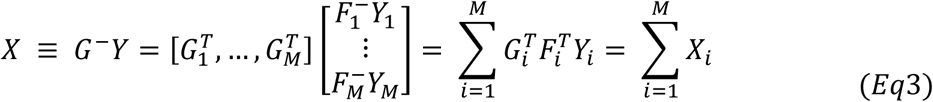

Here *M* is the number of subjects, and *G*^−^ is the *T*_2_by *MT*_1_standardized reducing matrix. Group-level principal components that explained the maximum variance were selected as the input for spatial ICA to calculate independent group components. We express the relationship as:

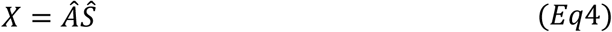

where the generative linear latent variables *Â* and *Ŝ* are the *T*_2_ × *T*_2_ mixing matrix related to subject time courses and the *T*_2_ × *V* aggregate spatial maps, respectively. In our proposed approach we modified (*Eq4*) as

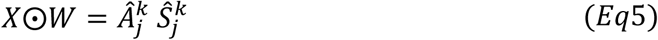

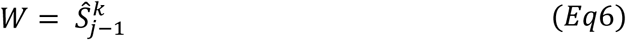

Where 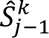 represent *k*^*th*^ spatial map from (*j* − 1)^*th*^ scale, and ⊙ is vector outer product. In the first scale, we select *W* = 1 as there is no previous scale (*j* = 0). In our proposed approach, group spatial ICA (GICA) was applied on the weighted signals (voxel wise multiplication of a network on the BOLD signal), leading to the identification of the networks at the next scale. (Figure 1)

**Figure 1:**
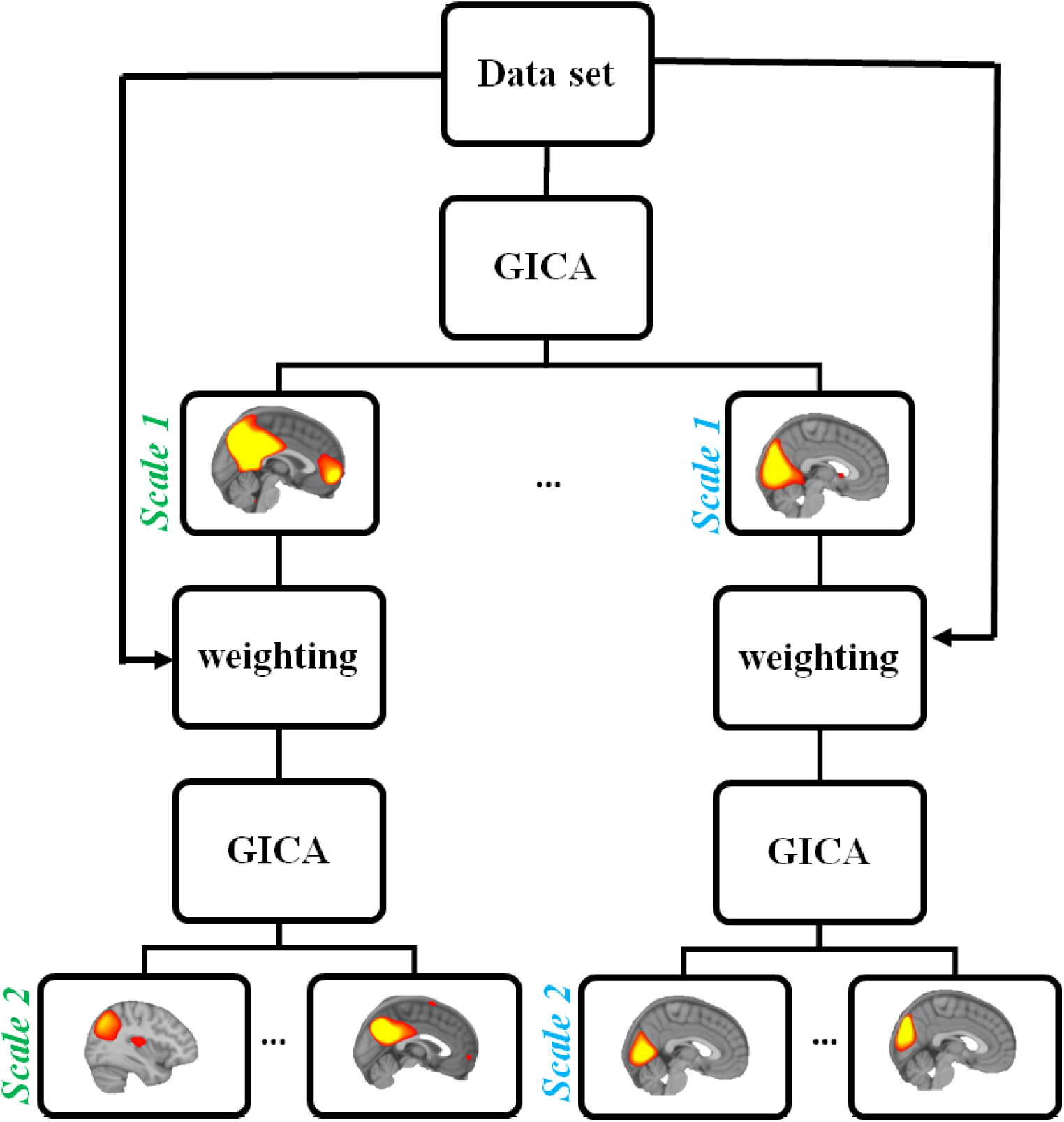
Introducing Telescopic independent component analysis (TICA): a novel approach for identifying Intrinsic connectivity networks (ICNs) at the multis-scale level. In each scale group-level spatial ICA(GICA) was applied on the weighted signal (voxel-wise multiplication of a network on the BOLD signal), leading to the identification of the network at the next scale. Since there is no previous scale in the first scale, GICA is applied to the original BOLD signal.

### 2.2. Demographics and data acquisition

We utilized a data set from the Functional Imaging Biomedical Informatics Research Network (FBIRN) for our investigation (Potkin & Ford, 2009). The FBIRN dataset was collected from seven sites. The same rs-fMRI parameters were used across all sites: a standard gradient echo-planar imaging (EPI) sequence, repetition time (TR)/echo time (TE) = 2,000/30 ms, voxel spacing size = 3.4375 × 3.4375 × 4 mm, slice gap = 1 mm, flip angle (FA) = 77°, field of view (FOV) = 220 × 220 mm, and a total of 162 volume. Six of the seven sites used 3-Tesla Siemens Tim Trio scanners, and one used a 3-Tesla General Electric Discovery MR750 scanner. The dataset was chosen using the following quality control: (a) individuals diagnosed with either typical control or schizophrenia conditions, (b) data demonstrating high-quality registration to the EPI template, (c) minimal head motion characterized by less than 3° rotations and 3-mm translations in every direction, and (d) mean framewise displacement less than 0.2. Consequently, our analysis was performed on rs-fMRI data derived from 310 individuals, comprising 150 HC and 160 SZ, as summarized in Table 1. All participants are also at least 18 years old and gave written informed consent before enrollment.

**Table 1:** Demographic information of the data used in the Functional Imaging Biomedical Informatics Research Network (FBIRN) study.

| FBIRN | Diagnostic | Numbers | sex | Numbers | <u>Age(years)</u> |  |
| --- | --- | --- | --- | --- | --- | --- |
| | | | | | Mean ± STD | |
|  |  |  |  |  | Median/Range |  |
| FBIRN | Healthy group | 160 | Male | 115 | 37±10.71 | 39(19/59) |
| | | | Female | 45 | 36.47±11.33 | 33(19/58) |
| | Schizophrenia group | 150 | Male | 114 | 38.74±11.78 | 40(18/62) |
| | | | Female | 36 | 39.06±11.40 | 36(21/57) |

### 2.3. Analysis pipeline

The re-fMRI data preprocessing was conducted using the Statistical Parametric Mapping toolbox (SPM12, https://www.fil.ion.ucl.ac.uk/spm/). Preprocessing steps include discarding the initial five volumes, rigid motion correction, and slice-time correction. Subsequently, individual subject data were registered to an MNI EPI template, resampled, and smoothed with a 6mm Gaussian Kernal, and variance normalization was applied on voxel time (Armin Iraji et al., 2022). We used the GIFT toolbox https://trendscenter.org/software/gift/ (Calhoun & Adalı, 2012; Calhoun et al., 2001; Iraji et al., 2021) to apply TICA. To begin, the 30 PCs that explained the maximum variance of each subject and the 20 group-level PCs that explained the maximum variance of each estimator-specific data set were used as the input for spatial GICA. We selected a spatial GICA model order of 20 to obtain large-scale ICNs (Iraji et al., 2016; Iraji et al., 2019). For the GICA algorithm, we used the infomax algorithm (Bell & Sejnowski, 1995; Lee et al., 1999) ran it 100 times, and utilized the ICASSO framework to identify the best estimate, defined as the independent component closest to the stable cluster (Himberg et al., 2004). We employed back-reconstruction using a GICA technique (Calhoun et al., 2001) to compute subject-specific independent components and time courses because it provides more accurate time courses and spatial maps.

Afterward, we employed a selection process for identifying ICNs. We identified ICNs using prior knowledge and the following criteria (a) dominant low-frequency fluctuations in their time courses, determined through dynamic rage assessment and the low-frequency to high-frequency power ratio;(b) peak activation located within gray matter regions;(c) minimal spatial overlap with vascular and ventricular structures and (d) low spatial similarity with motion and other recognized artifacts (E. Allen et al., 2011). Next, we picked DMN, VS, and RFPN and performed the weighting step for all subjects’ BOLD signals with their spatial map of scale 1. Finally, we applied the GICA to the weighted BOLD signals. Similar to the previous scale, we applied 2 levels of PCA. In subject level 6 PCs, and group level 4 PCs that explained the maximum variance of the data set were used to input for GICA. Previous works(Abou-Elseoud et al., 2010; E. A. Allen et al., 2011; Hu et al., 2016; A. Iraji et al., 2022; Li et al., 2007) highlight the impact of model order on ICA results; we used elbow point criteria to determine the optimal number of PCs. All other parameters in GICA to extract ICNs scale 2 are the same as scale 1. We also applied GICA scale 2 with model orders 2,3,5, and 6 on DMN to investigate how model order choice affects the results of TICA.

### 2.4. Accessing the replicability of the TICA approach-pipeline evaluation

We evaluated the replicability (Adali et al., 2022; National Academies of Sciences et al., 2019) of our approach using the initial random division of the dataset into two distinct subsets, each of which underwent independent processing through our pipeline. This division and processing were repeated 100 times, with each iteration yielding one DMN scale 1 and its corresponding four zoomed-in DMN, extracted from scale 2. To quantify the replicability of our approach, we calculated the average of spatial similarity between each pair of ICN from two independent half splits using the Pearson correlation (Figure 2).

**Figure 2:**
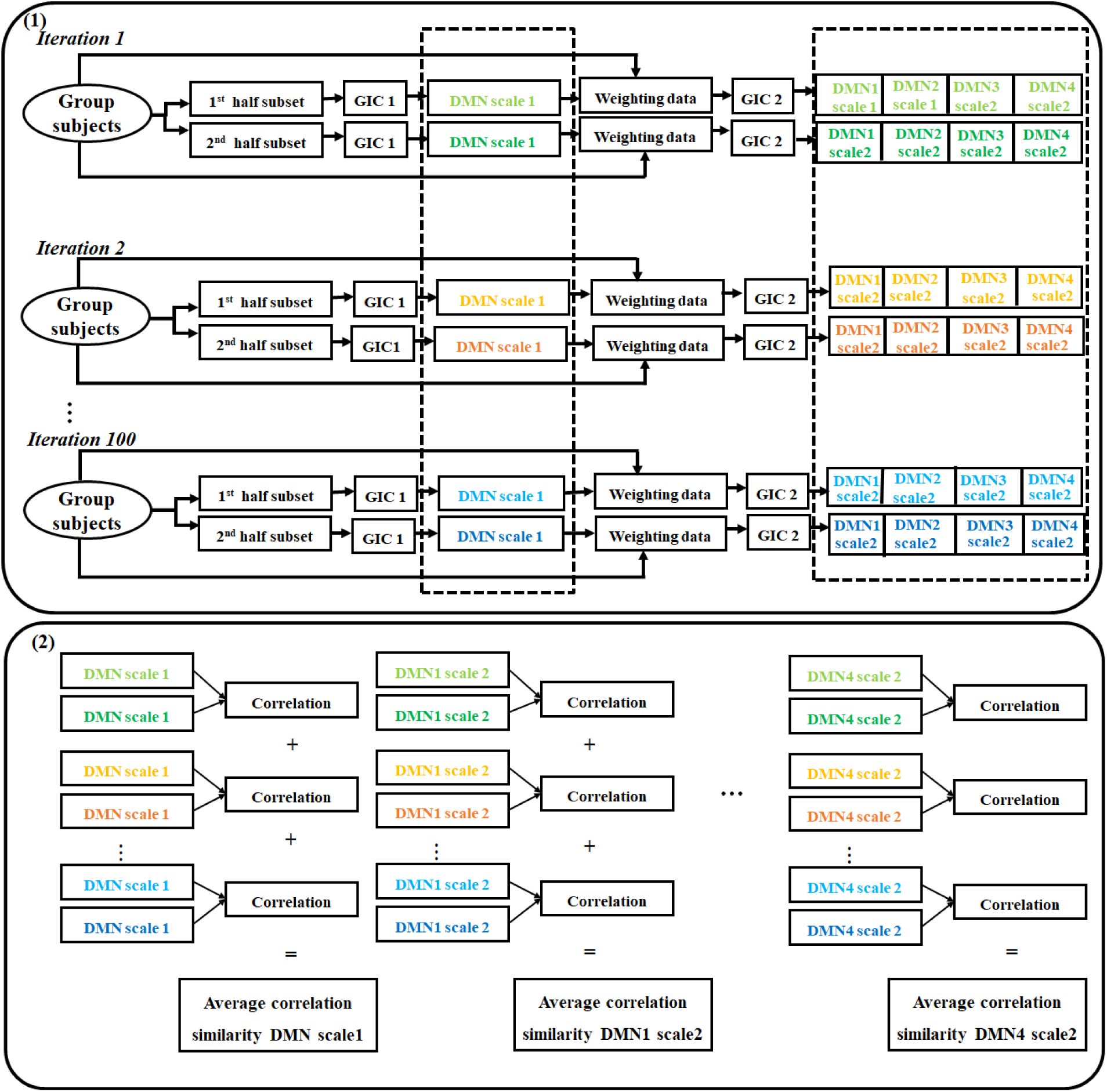
Pipeline to assess the replicability of the TICA approach. First, the dataset was divided into two separate subsets randomly. Each of these subsets was subjected to independent processing through the TICA approach. This division and processing procedure were repeated 100 times, leading to the extraction of the DMN from scale 1 and its corresponding four zoomed-in DMN obtained from scale 2. Finally, the average spatial similarity was calculated by using the Pearson correlation for every pair of networks within the two subsets.

We also assessed the similarity between individual-level DMN and group-level DMN in scale 1 and scale 2; as well as the similarity between individual-level DMN in scale 1 and scale 2. To accomplish this, we obtained the DMN for each participant; and we computed the Pearson correlation between the group-level DMN and corresponding individual DMN to get the similarity in group and individual level. We also calculated the Pearson correlation between individual subjects and obtained the similarity at the individual level. This comparative analysis was performed for both scale 1 and scale 2 DMN.

Moreover, we conducted additional analysis to assess the test-retest reliability on TICA to specifically measure individual differences. We used the Human Connectome Project (HCP) (Van Essen et al., 2013)data sets. The HCP data set consists of 1019 subjects and has four complete rsfMRI scan sessions. Each session has 1200 time points, a TR = 0.72 sec, and an original voxel size of 2.0 mm isotropic. We ran TICA separately on two different scan sessions and calculated the Pearson correlation similarity between group-level DMNs in scale 1 and scale 2. We also calculated the Pearson correlation similarity for each subject between two different scan sessions.

### 2.5. Group comparison analysis

To assess ICN differences between HC and SZ, we first identified the voxels with a strong contribution to a given ICA (Z>1.96). We next conducted voxel-wise two-sample t-tests and applied an adaptive thresholding technique based on the Gamma-Gaussian mixture model (Gorgolewski et al., 2012)to identify voxels with significant diagnostic effects.

Moreover, to evaluate the consistency of group differences, we calculated the group difference using a random 80% subset of the dataset. We then measured the similarity between the T-map generated from the entire dataset and the T-map from this 80% subset. This process was repeated 50 times.

### 2.6. High order GICA

To investigate the comparison of DMN components derived through the TICA method using a single scale ICA, we conducted ICA with a high model order on the FBIRN data set. The GICA analysis steps are as follows. First, 120 PCs that explained the maximum variance of each subject were retained for further analysis. Next, 80 group-level PCs that explained the maximum variance of each estimator-specific data set were used as the input for GICA. We applied model order 80 as GICA high order; The model order choice was motivated by the TICA procedure, where we initially employed GICA with a model order of 20 to get networks in scale 1, then an additional GICA with a model order of 4 used to get smaller networks in scale 2. For the high model order GICA algorithm, similar to the TICA approach, we used the infomax algorithm, the ICASSO framework, and GICA back-reconstruction techniques to compute subject-specific independent components and time courses. Afterward, we employed a selection process for identifying the DMN like the TICA approach.

To quantify the comparing the results between TICA and high-order GICA, we calculated two different approaches. First, we evaluated the similarity between spatial maps generated by high order ICA and TICA approach using Pearson correlation. Second, we conducted a paired-sample t-test on spatial maps to compare the TICA approach and high model order ICA across DMNs.

We also applied group comparison analysis for DMN derived from high-order GICA. Similar to the TICA approach, we identified the group effect in voxels that significantly contribute to DMN in HC and SZ groups using an adaptive thresholding technique to identify regions with significant diagnostic effects. Moreover, to evaluate the consistency of group differences, we calculated the group difference using a random 80% subset of the dataset. We then measured the similarity between the T-map generated from the entire dataset and the T-map from this 80% subset. This process was repeated 50 times to find similarity between the T-maps for all components.

## 3. Results

### 3.1. The intrinsic connectivity network hierarchy

According to the criteria specified in the materials and methods section, we identified DMN, VS, and RFPN extracted from scale 1, along with four ‘zoomed in’ ICNs derived from scale 2. The decomposition of the ICNs across these two spatial scales is visually depicted in (Figure 3).

**Figure 3:**
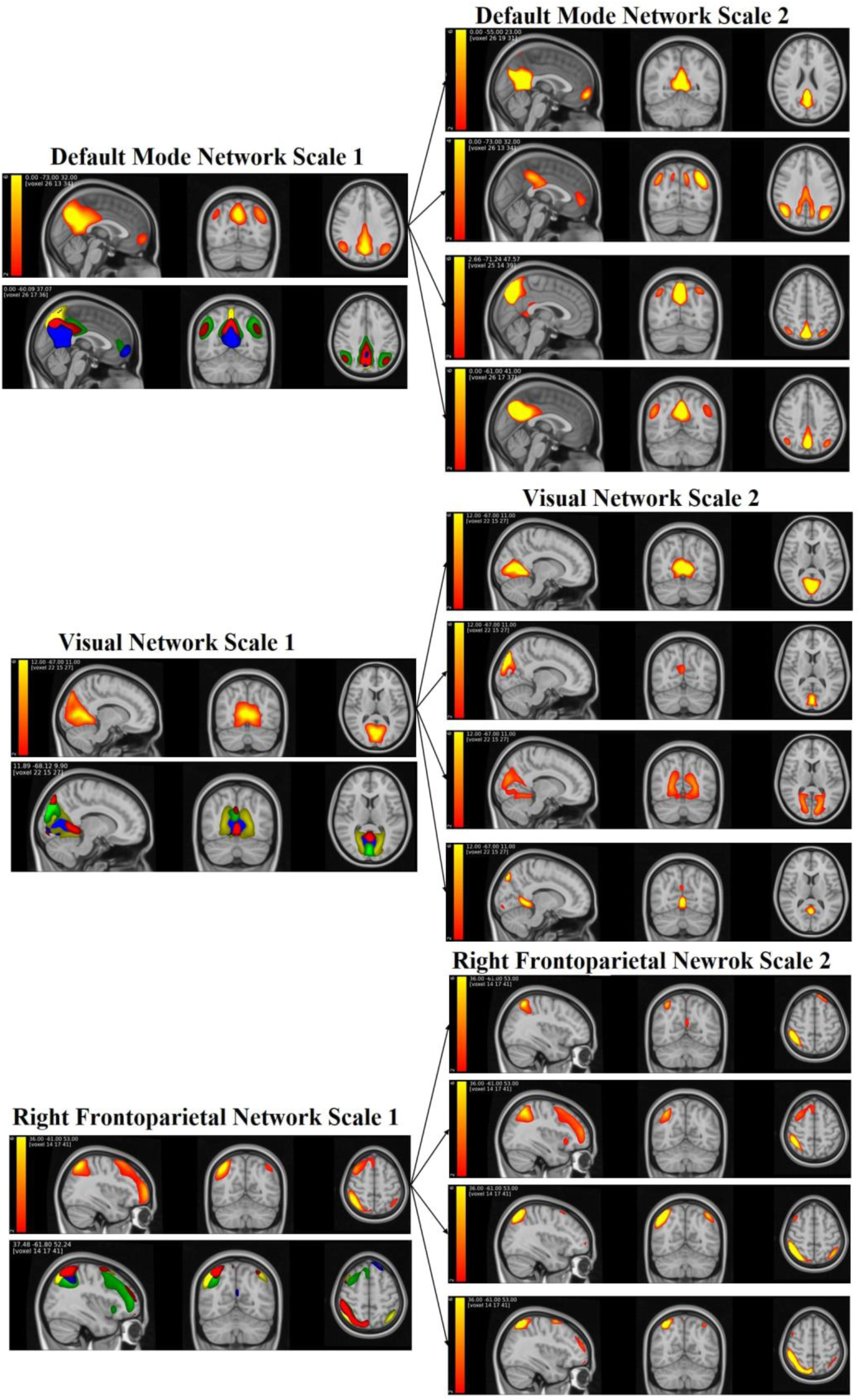
Visualization of DMN, VN, and RDPN spatial maps extracted from the TICA approach. It shows the decomposition of the three networks scale 1 in scale 2.Blue: first component in scale 2, green: second component in scale 2, yellow: third component in scale 2, and red: fourth component in scale 2

In DMN, VS, and RFPN scale 1, the number of voxels that significantly contribute (Z > 1.96) in spatial maps is 2948, 3122, and 3307 respectively. In scale 2, the DMN significant voxels are 1077, 2723, 965, and 1426 respectively. VS significant voxels are 1544, 1239, 2860, and 1058 respectively. Significant voxels in RFPN spatial maps are 1389, 2777, 968, and 1130 respectively. These shows that the TICA approach captures bigger networks in scale 1 and smaller networks in scale 2. In addition, the spatial correlation similarities between DMN scale1 with DMN scale2 are 0.43, 0.61, 0.42, and 0.61 respectively. The similarity in VS between scale 1 and scale 2 are 0.72, 0.47, 0.49, and 0.25. The similarity in RFPN between scale 1 and scale 2 are 0.27, 0.70, 0.54, and 0.26 respectively. These results display that the telescopic approach can capture distinctive networks in each scale.

Figure 4 (A) displays the Pearson correlation coefficient between the DMN in scale 1 and scale 2 across the first and second halves of the data set over 100 iterations. The replicability index (average spatial similarity across 100 iterations) for all ICNs is well above 0.8. The DMN scale 1 exhibited a reliability of 0.97 with a standard deviation of 0.01. The DMN generated in scale 2 also yielded replicability with values of 0.80, 0.83, 0.91, and 0.81 along with corresponding standard deviations of 0.20, 0.18, 0.06, and 0.24, respectively. The lower average similarity in scale 2 compared with scale 1 indicated some variability in more detailed functional sources. This outcome aligns with the observed pattern in a high model order ICA approach (Abou-Elseoud et al., 2010).

**Figure 4:**
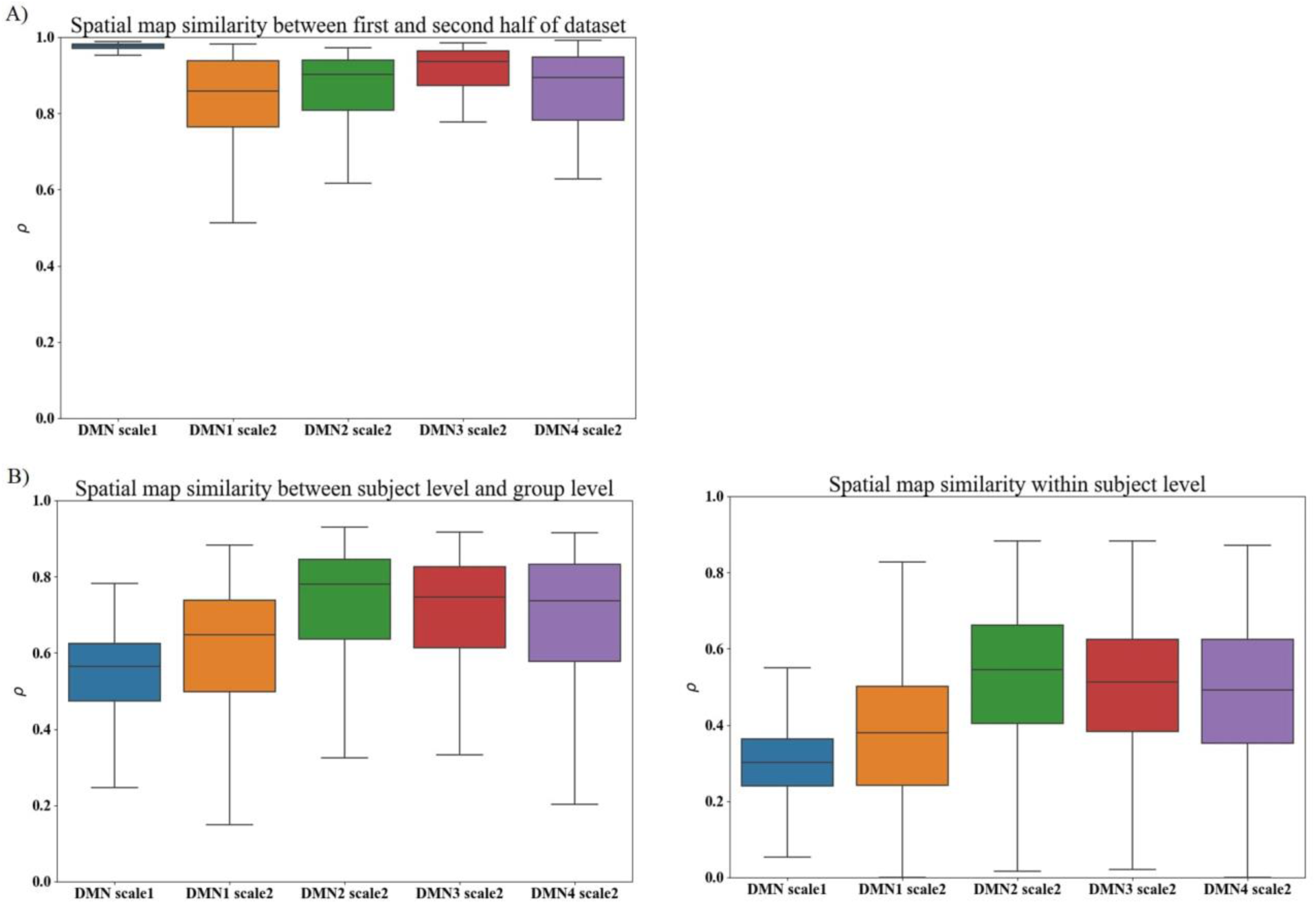
(A) Pearson correlation coefficient between the first and second half of the dataset, providing insights into the pipeline’s replicability. The average similarities of all components are higher than 0.8. (B) Pearson correlation coefficient between subject-level and group-level. The average similarity in scale 2 is higher than scale 1 in both case comparisons.

We also present the similarity between individual-level DMN spatial maps as well as individual-level and group-level DMN spatial map generated from the TICA approach using the Pearson correlation coefficient. The computed average similarities between group-level and individual-level on scale 1 was 0.54, and on scale 2 were 0.60, 0.72, 0.70, and 0.69, respectively. Also, the average Pearson correlation coefficient in individual-level from DMN scale 1 and DMN scale 2 are 0.30, 0.37, 0.52, 0.49, 0.48. Standard deviations of correlation similarity in individual-level are 0.09, 0.17, 0.18, 0.17, and 0.18 respectively. The visual representation in Figure 4 (B) illustrates the boxplot of these similarities, revealing a higher similarity in DMN scale 2 as compared to DMN scale 1.

(Figure 5) displays the Pearson correlation coefficient for each subject between two scan sessions evaluating the test-retest reliability of TICA approach. The results show that the correlation coefficients in scale 2 are higher than in scale 1. In addition, the correlation similarities of group-level DMNs between two scan sessions in scale 1 is 0.98 and scale 2 are, 0.99, 0.99, 0.97, and 0.99 respectively.

**Figure 5:**
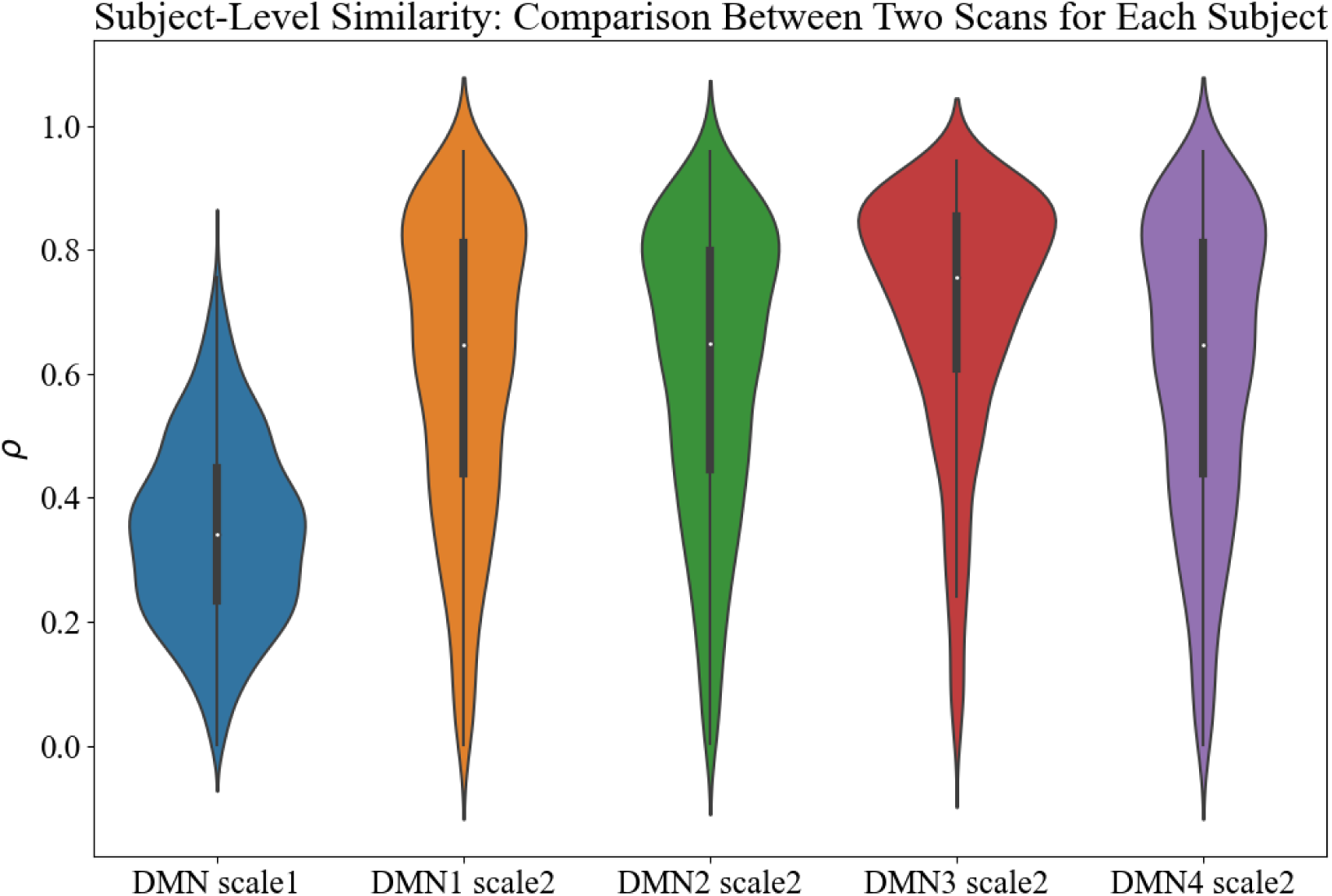
Pearson correlation similarity of subject level DMNs between two scan sessions. The similarity in scale 2 is higher than scale 1.

### 3.2. Model order effect

To study how model order choice affects the TICA results, we conducted analyses on TICA scale 2 with model orders 2,3,5, and 6. In general, results (Figure 6) are similar and consistent with the previous ICA work. As model order increases the components branch into multiple components (Abou-Elseoud et al., 2010); for instance, the first component in order 2 is formed by combining the first and second components from order 4. The finding also highlights that model orders 3-5 showed spatial overlaps of components.

**Figure 6:**
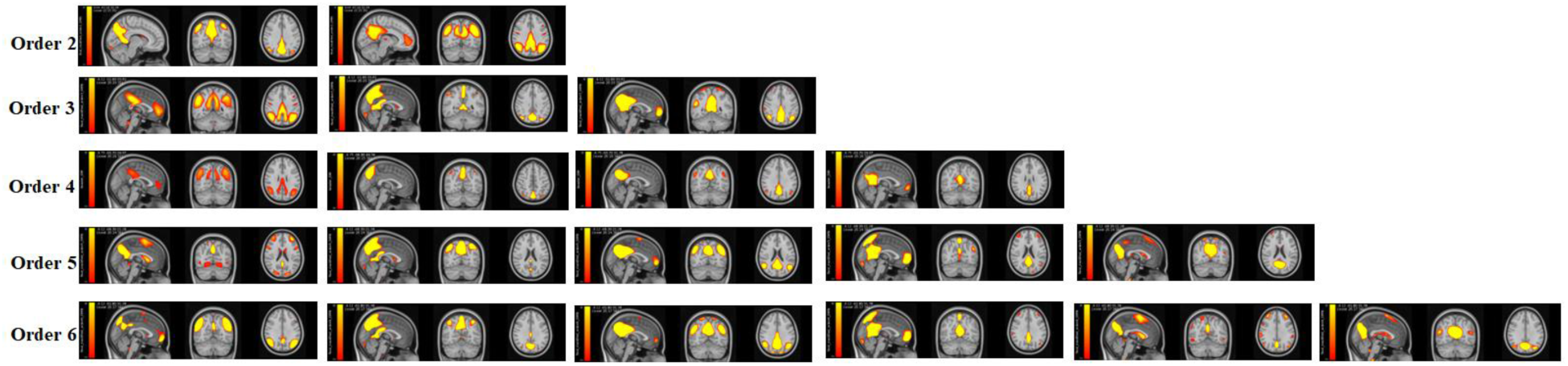
Spatial maps generated by the TICA approach with model orders 2,3,4,5, and 6 in scale 2. The results show the model order impacts the quality of the results.

### 3.3. Clinical applications

(Figure 7) illustrates the t-map of voxels with significant diagnostic effects (*p* < 0.01) after adoptive thresholding using Gamma-Gaussian mixture model. Notably, the maps revealed significant group differences in the DMN scale2, which were not evident in scale1. In DMN scale 1, the areas exhibiting significant differences include the middle cingulate and paracingulate gyri, posterior cingulate gyrus, cuneus, lingual gyrus, middle occipital gyrus, angular gyrus, and precuneus. Conversely, in DMN scale 2, significant differences besides the area in scale 1 are manifested in the superior frontal gyrus, medial orbital, calcarine fissure, surrounding cortex, superior occipital gyrus, and inferior parietal gyrus.

**Figure 7:**
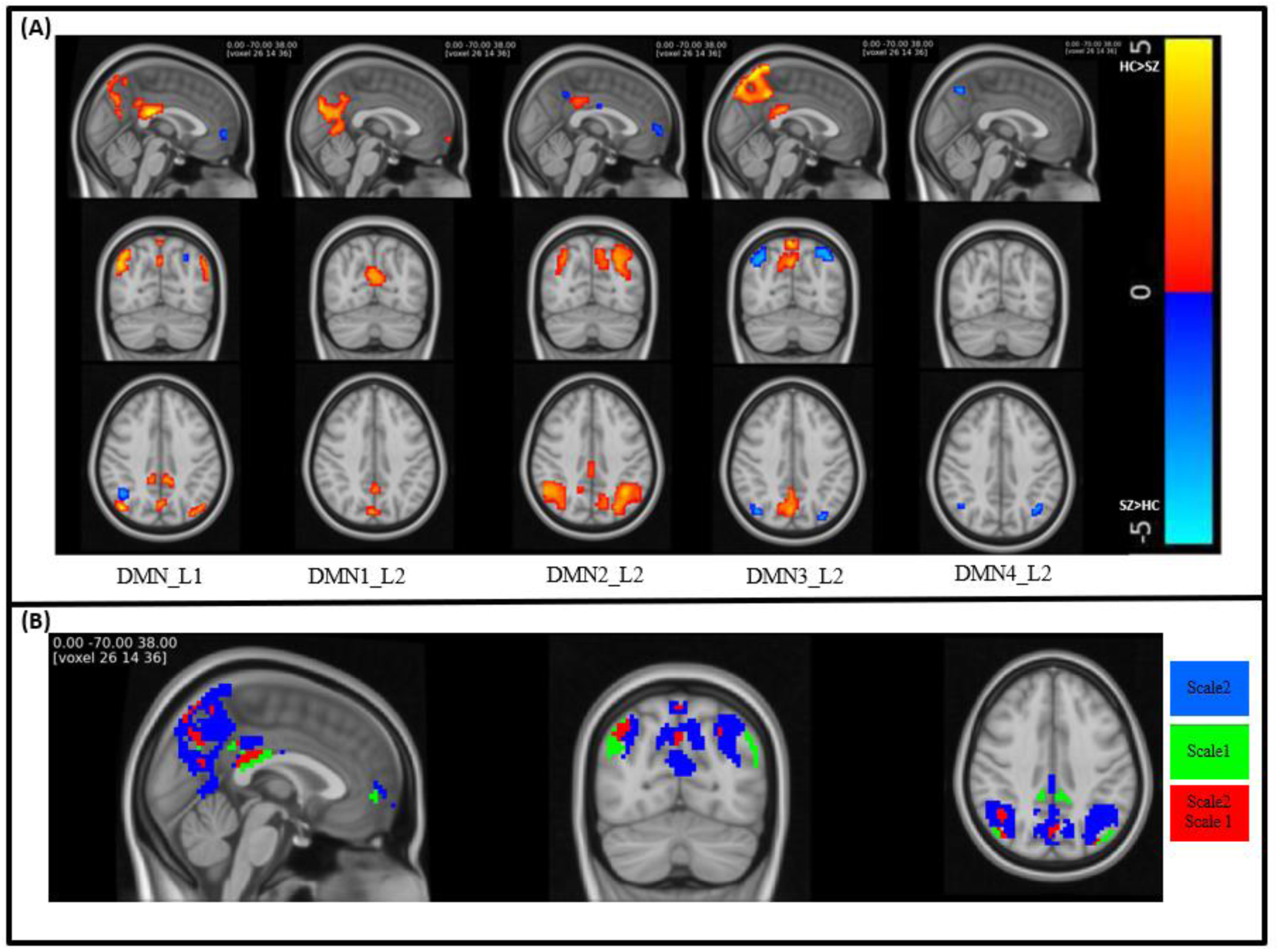
(A)visualization of T-map of brain regions show significant group differences (*p* < 0.01) between HC and SZ under TICA approach. (B)Visualization of significant voxels in both scale 1 and scale 2(red), only scale 1(green), and only scale 2(blue).

(Figure 8) shows consistency in group differences uing Pearson correlation coefficient between T-maps generated from the group difference across the entire dataset and 80% of the datasets over 50 iterations. The results revealed that the average similarity exceeded 0.90 for each component illustrating the consistency in group differences.

**Figure 8:**
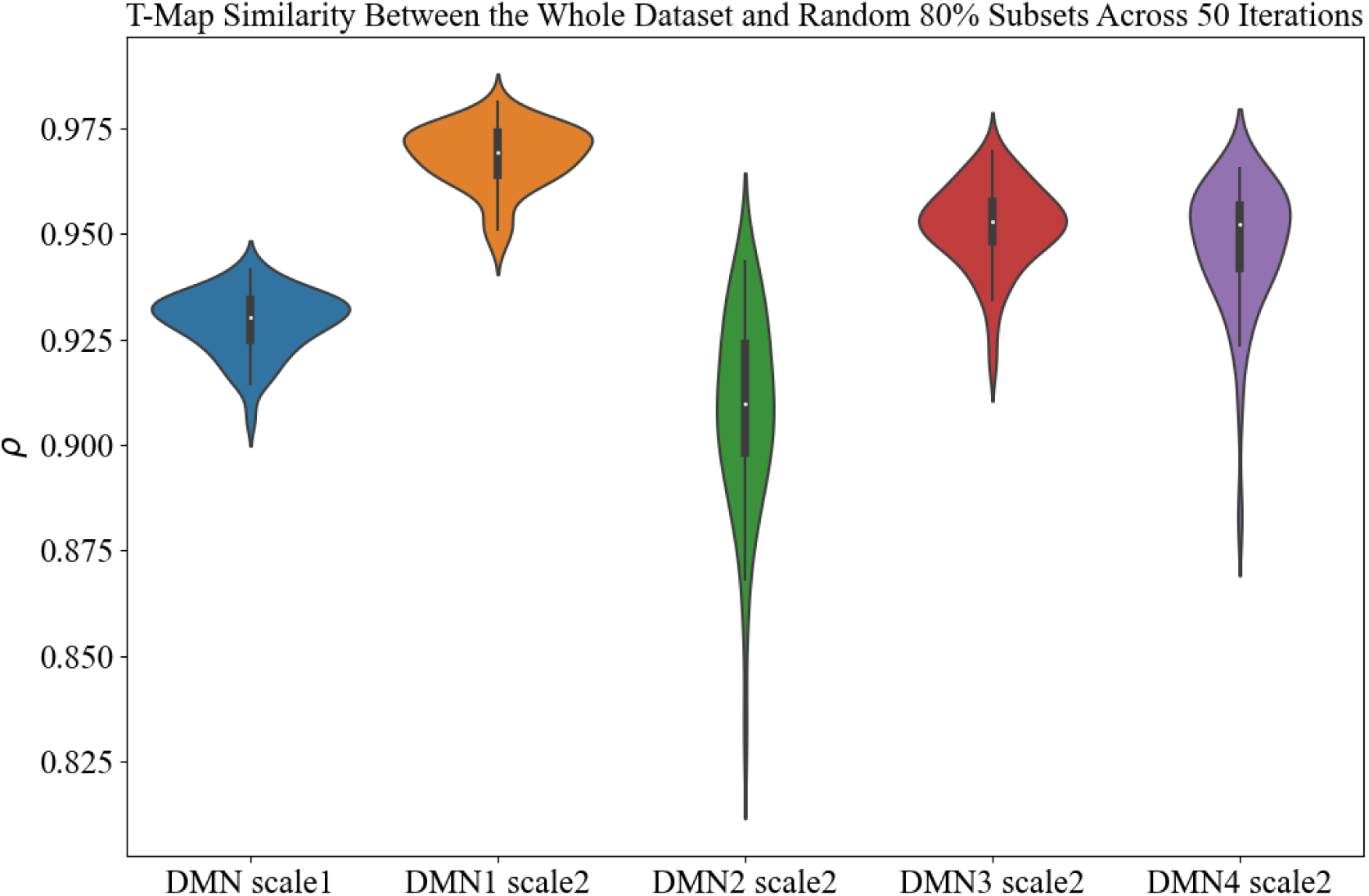
Consistency in group differences assessment: Pearson correlation between T-maps generated from group differences across the entire dataset and 80% of the datasets over 50 iterations. The average similarity exceeds 0.90 for each component illustrating the consistency in group differences.

### 3.4. GICA high order

We identified four distinct DMN extracted through ICA at order 80. (Figure 9. A) displays the spatial maps of these DMNs, denoted as DMN1, DMN2, DMN3, and DMN4. Additionally, (Figure 9. B) presents the outcomes of significant group differences between a HC and SZ, derived from all DMN at a high model order. The identified areas exhibiting significant group differences encompass the inferior parietal gyrus, angular gyrus, precuneus, and cuneus regions. The number of significant voxels in DMN from ICA order 80 are revealed as 49, 46, 32, and 14, respectively.

**Figure 9:**
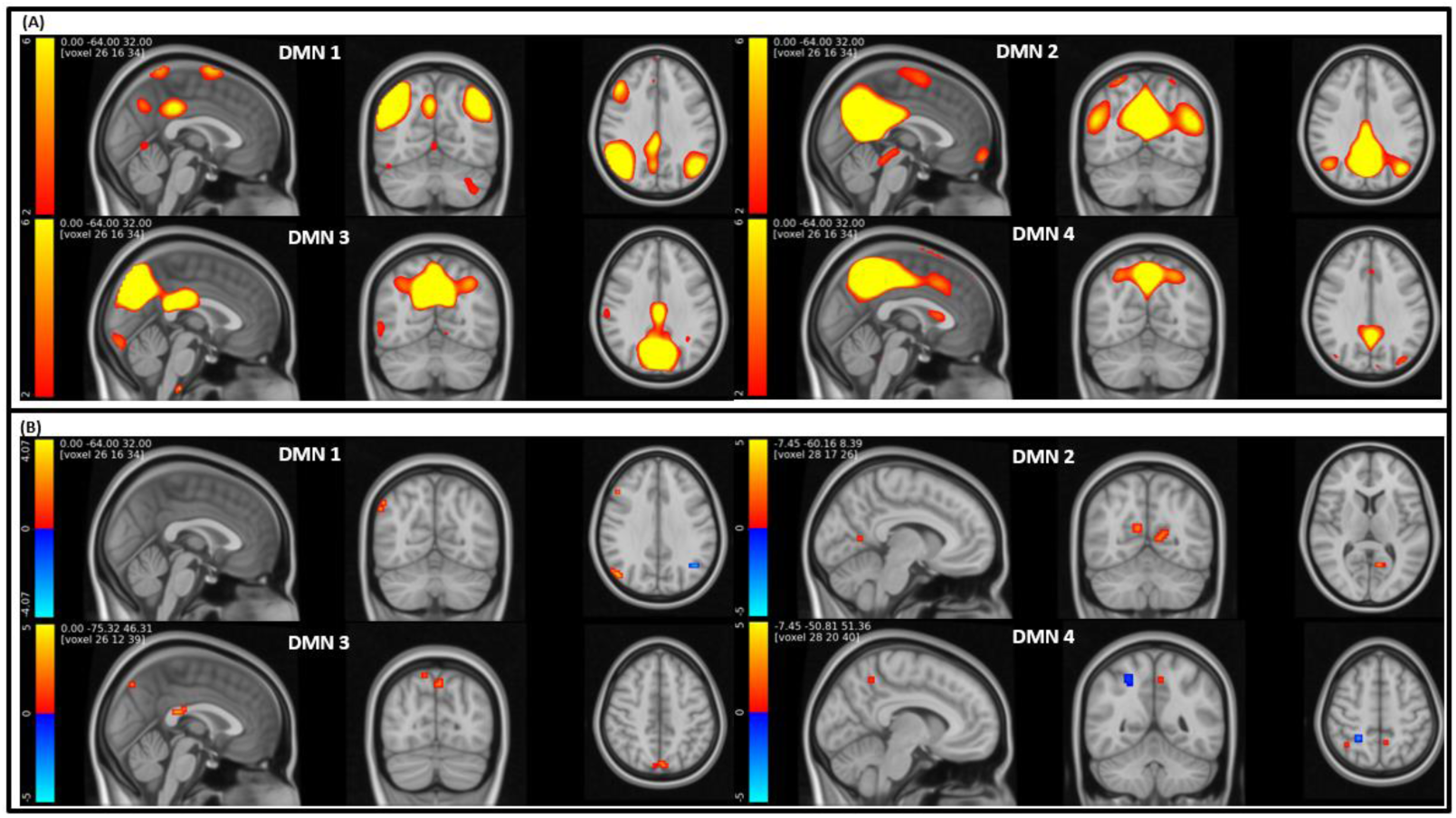
(A)Visualization of DMN spatial maps extracted from ICA order 80. (B) brain regions with significant group difference between HC and SZ in DMN area extracted from ICA order 80 after adoptive thresholding technique (p<0.01)

### 3.5 Comparison between TICA and high model order approaches

We conducted a paired-sample t-test to compare TICA and high model order approaches across DMNs. (Figure 10) displays voxels that show significant differences (P-value <0.05) after FDR correction between the two approaches. In addition, we assessed the correlation similarity between DMN1 high model order ICA and DMN generated from a telescopic approach, revealing values of (0.30, 0.13, 0.35, 0.10, 0.18). Similar correlation assessments were conducted for DMN2 high model order ICA (0.71, 0.57, 0.15, 0.19, 0.73), DMN3 high model order ICA (0.40, 0.01, 0.007, 0.50, 0.44), and DMN4 high model order ICA (0.25, 0.10, 0.19, 0.04, 0.08).

**Figure 10:**
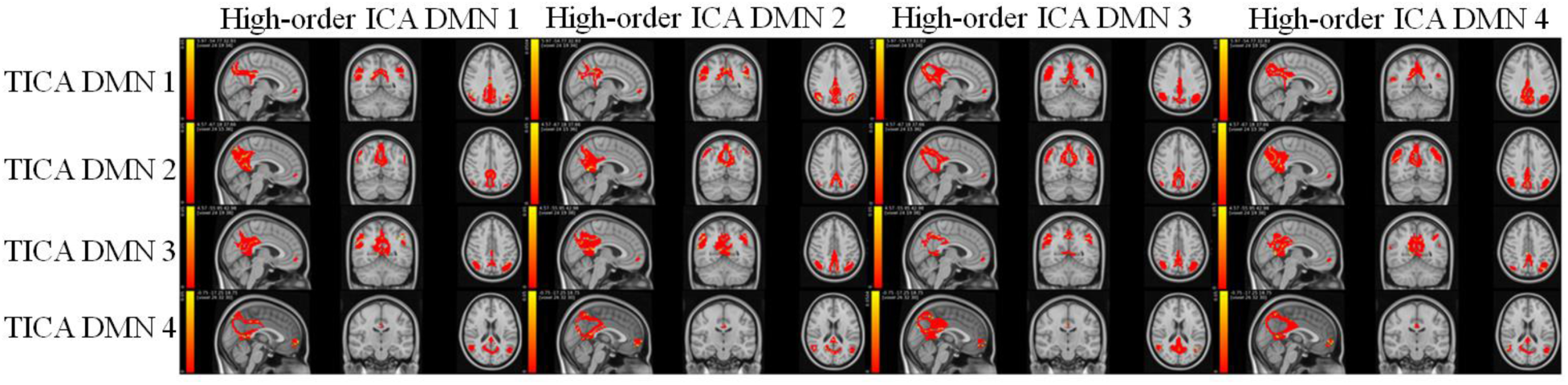
Comparison of DMN significant voxels identified by TICA and high model order approach. Voxels significantly different between the two approaches (P< 0.05) are highlighted.

## 4. Discussion

Studying brain functional connectivity has improved our understanding of brain functions and the impact of brain disorders. However, current studies in functional connectivity disregards functional connectivity across multiple spatial scales. At best, apply multimodal ICA (Armin Iraji et al., 2022)to study functional interactions. Still, they do not construct the spatial hierarchy or impose direct constraints on the size of ICNs. In this work, we introduced TICA as an effective and adaptive tool to identify ICNs construct spatial functional hierarchies and estimate functional sources across multiple spatial. We also leveraged the approach to study group comparison between HC and SZ which has been understudied as most research focuses on only a single spatial scale.

Using TICA, we identified five distinct DMNs, including the original network extracted from DMN scale1 and its decomposition in four small networks from DMN scale2. (3.1) supports the finding that how the TICA approach captures bigger the network in the first scale and uses its information to extract smaller networks in the next scale.

We observed that DMN in the first scale demonstrates a commendable mean reliability of 0.97, highlighting its robustness in effectively detecting the overall DMN. Additionally, the networks in the second scale exhibit reasonably high mean reliabilities, ranging from 0.80 to 0.91, indicating some variability in these more detailed functional sources. Our finding aligns with the observed pattern in a high model order ICA approach (Abou-Elseoud et al., 2010).

We observed a more pronounced group effect in Scale 2 in comparison to Scale 1 (3.2). Additionally, the results indicate that subject-level estimates exhibit greater similarity to both the reference and each other in Scale 2 as opposed to Scale 1 Figure 4 implying a more accurate estimation of networks. These collective findings suggest that the second scale may be more sensitive in capturing the group effect associated with SZ. Therefore, the TICA approach holds promise in identifying biomarkers for schizophrenia. We observed that DMN3 scale 2 resembles the spatial maps of the parietal memory network (PMN), a network with similar spatial patterns to the DMN but with distinct functional properties (Yang et al., 2014). This finding highlights that telescopic ICA can capture this distinct network and suggests it might be a network in the DMN hierarchy. Moreover, our analysis identifies significant group differences in the posterior cortex (PCC) and precuneus (PCU) area echoing similar observations in previous studies (Gilmore et al., 2015; Guo et al., 2020) that contribute to auditory hallucinations (AH) in schizophrenia. Additionally, while previous work estimated this network in the first layer and high-order ICA, our analysis suggests that leveraging telescopic ICA can improve our ability to differentiate between this network and others within the default mode domain.

Previous studies show higher model order ICA results in more spatially granular ICNs (E. A. Allen et al., 2011; A. Iraji et al., 2022); however, (3.4) show that the DMNs captured by the TICA approach are different from a single-scale ICA with high model order. Also, more significant group differences were captured using the TICA approach compared to high model order ICA. These results also emphasize the importance of studying functional brain networks on multi-scale using the TICA approach.

In this study, we focus on only two scales with ICA model order 20 and 4 to capture networks. Further studies can develop our approach in incremental scales to effectively capture ICNs in multiscale. Furthermore, future studies can explore differences across the different back-reconstruction approaches (Erhardt et al., 2011). Here, we also focused on the voxel-wise group comparison of DMN spatial maps. Future studies should focus on TICA approach using static functional network connectivity, and dynamic functional network connectivity on DMN as well as other brain networks(Armin Iraji et al., 2022).

## 5. Conclusion

Brain functional connectivity can occur at different spatial scales, which has been underappreciated. Here we proposed a TICA approach as a new tool to construct spatial functional hierarchies of brain and estimate functional sources across multiple spatial scales. Our results on DMN show that our approach can capture replicable comprehensive spatial maps. Moreover, the TICA can detect significant group differences between HC and SZ that are missed using studying brain networks in single scale highlighting that our proposed approach represents a promising new tool for studying functional sources.

## 6. Author contribution

Shiva Mirzaeian: Conceptualization; Methodology; Visualization; Writing-Original Draft. Ashkan Faghiri: Writing-Review & Editing. Vince Calhoun: Conceptualization; Investigation; Supervision; Writing-Review & Editing. Armin Iraji: Conceptualization; Investigation; Supervision; Writing-Review & Editing.

## Acknowledgements

Data collection was supported by the National Center for resources at the National Institute of Health [grant numbers: NIH 1 U24 RR021992, NIH 1 U24 RR025736-01].

